# Mechanically active integrins direct cytotoxic secretion at the immune synapse

**DOI:** 10.1101/2021.10.02.462778

**Authors:** Mitchell S. Wang, Yuesong Hu, Elisa E. Sanchez, Xihe Xie, Nathan H. Roy, Miguel de Jesus, Weiyang Jin, Joanne H. Lee, Yeonsun Hong, Minsoo Kim, Lance C. Kam, Khalid Salaita, Morgan Huse

## Abstract

The secretory output of cell-cell interfaces must be tightly controlled in space and time to ensure functional efficacy. This is particularly true for the cytotoxic immune synapse (IS), the stereotyped junction formed between a cytotoxic lymphocyte and the infected or transformed target cell it aims to destroy^1^. Cytotoxic lymphocytes kill their targets by channeling a mixture of granzyme proteases and the pore forming protein perforin directly into the IS^2, 3^. The synaptic secretion of these toxic molecules constrains their deleterious effects to the target cell alone, thereby protecting innocent bystander cells in the surrounding tissue from collateral damage. Despite the importance of this process for immune specificity, the molecular and cellular mechanisms that establish secretory sites within the IS remain poorly understood. Here, we identified an essential role for integrin mechanotransduction in cytotoxic secretion using a combination of single cell biophysical measurements, ligand micropatterning, and functional assays. Upon ligand-binding, the α_L_β_2_ integrin LFA-1 functioned as a spatial cue, attracting lytic granules containing perforin and granzyme and inducing their fusion at closely adjacent sites within the synaptic membrane. LFA-1 molecules were subjected to pulling forces within these secretory domains, and genetic or pharmacological suppression of these forces abrogated cytotoxicity. We conclude that lymphocytes employ an integrin-dependent mechanical checkpoint to enhance both the potency and the security of their cytotoxic output.

---

Lytic granules, the specialized secretory lysosomes that store perforin and granzyme, are known to accumulate at the IS and fuse selectively with the synaptic membrane^2, 3^. This behavior has long been attributed to the centrosome, which serves as a focal point for intracellular granules and polarizes toward the target cell during IS formation^2, 4^. Studies from our group and others, however, indicate that the centrosome is dispensable for synaptic secretion, implying the existence of other targeting mechanisms^5-7^. Cytotoxic lymphocytes exert nanonewton scale forces across the IS that have been implicated in both the activation of mechanosensitive cell surface receptors and the potentiation of perforin function^8-14^. We have found that lytic granule exocytosis (degranulation) tends to occur in regions of active force exertion within the IS^8^, raising the possibility that local mechanosensing might play an instructive role in guiding perforin and granzyme release.

To explore this hypothesis, we studied the degranulation requirements of primary murine CD8^+^ cytotoxic T lymphocytes (CTLs), which are activated by T cell receptor (TCR)-mediated recognition of antigenic peptide-major histocompatibility complex (pMHC) on the target cell. The CTLs we used for our experiments expressed the OT-1 TCR, which is specific for the ovalbumin_257-264_ peptide (OVA) bound to the class I MHC protein H2-K^b^. pMHC recognition by the TCR triggers IS formation and the “inside-out” activation of cell surface integrins^1, 15^, including LFA-1 (Lymphocyte function-associated antigen-1), which establishes strong adhesion by binding to its cognate ligands Intercellular adhesion molecule-1 (ICAM-1) and ICAM-2 on the opposing membrane. Using micropillar-based traction force microscopy^16^, we found that LFA-1 engagement substantially increased synaptic force exertion (Fig. 1a-b), implying that this integrin is a key player in IS mechanics and therefore a candidate for mediating coupling between mechanical and secretory output.

**Figure 1.**
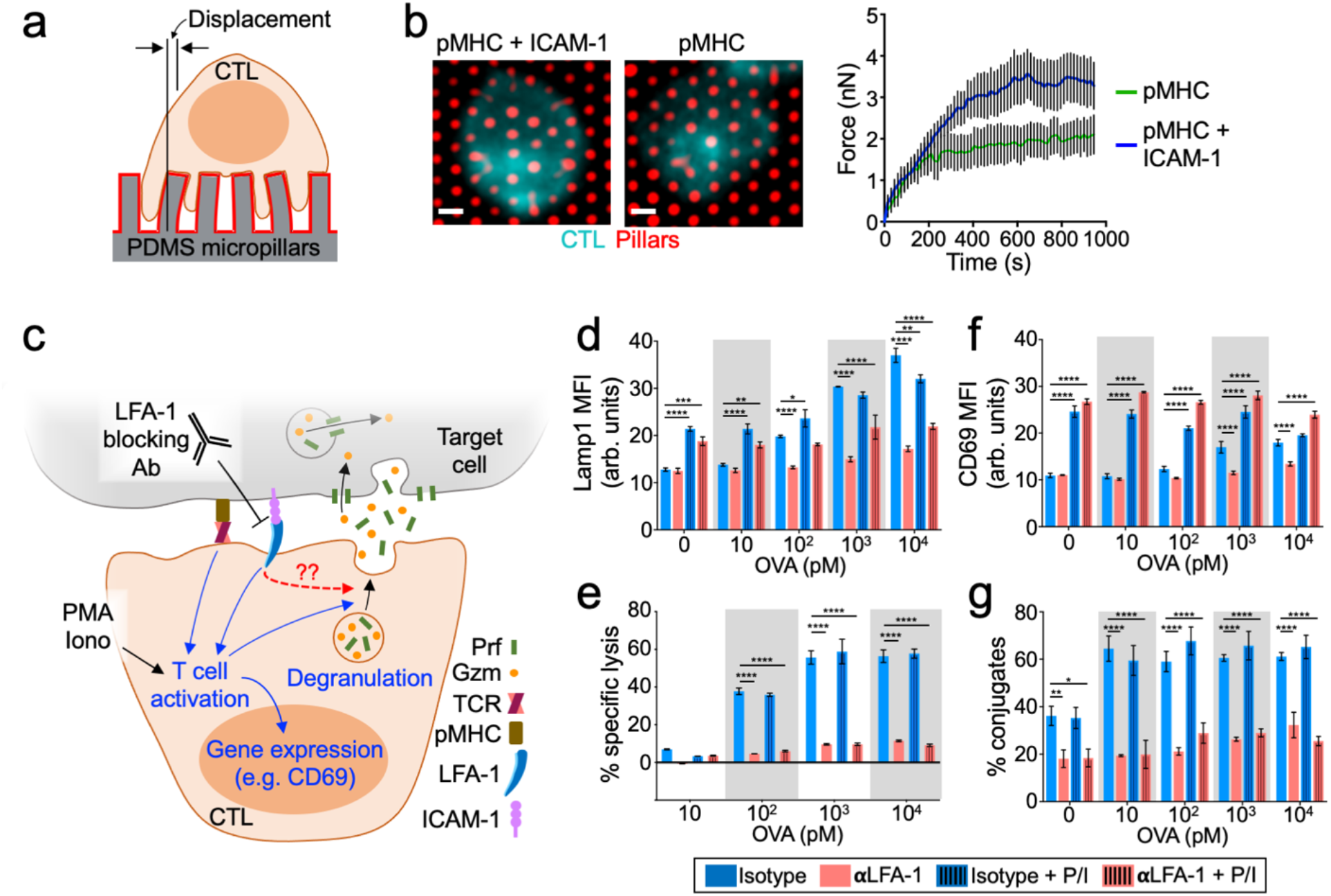
LFA-1 is required for synaptic force exertion, degranulation, and cytotoxicity. (a) OT-1 CTLs labeled with fluorescent anti-CD45 Fab were stimulated on PDMS micropillar arrays coated with pMHC ± ICAM-1. Traction forces were derived from pillar displacement. (b) Left, representative images of CTLs interacting with the arrays. Scale bars = 2 μm. Right, mean force exertion against the array graphed against time. N ≥ 8 for each sample. (c) OVA-loaded RMA-s target cells were mixed with OT-1 CTLs in the presence of LFA1 blocking antibody (αLFA-1) or isotype control. PMA/Iono was applied to some samples in order to drive TCR independent CTL activation. (d) Lamp1 exposure (degranulation), measured 90 min after CTL-target cell mixing. (e) Target cell killing, measured 4 h after CTL-target cell mixing. (f) CD69 expression, measured 90 min after CTL-target cell mixing. (g) Conjugate formation, measured 90 min after CTL-target cell mixing. Data in d-g were derived from technical triplicates. All error bars denote SEM. *, **, ***, and **** denote P ≤ 0.05, P ≤ 0.01, P ≤ 0.001, and P ≤ 0.0001, calculated by 2way ANOVA. All data are representative of at least two independent experiments.

To evaluate the importance of LFA-1 for degranulation and cytotoxicity, we treated cocultures containing OT-1 CTLs and OVA-loaded RMA-s target cells with a neutralizing antibody that specifically disrupts LFA-1-ICAM-1/2 binding (Fig. 1c). Blocking LFA-1 in this manner dramatically inhibited antigen-induced degranulation, which we measured by surface exposure of the lysosomal marker Lamp1 (Fig. 1d and Fig. S1a) and by the depletion of intracellular granzyme B (Fig. S1b). Anti-LFA-1 neutralizing antibodies also strongly suppressed cytotoxicity, as quantified by lysis of target cells (Fig. 1e). LFA-1 engagement was similarly important for degranulation responses elicited by stimulatory beads coated with pMHC and/or ICAM-1 (Fig. S1c-e). Collectively, these results were consistent with a specific role for LFA-1 in degranulation, but they did not rule out the possibility that LFA-1 engagement might promote this response secondarily by augmenting T cell activation (Fig. 1c). Indeed, LFA-1 is known to function as a “costimulatory” receptor by lowering the antigen threshold required for signaling and TCR-induced gene expression^17, 18^. In our hands, LFA-1 blockade altered some, but not all, of these responses. We observed no effect on antigen-induced proliferation (Fig. S2a), assessed by dilution of CellTrace Violet dye. Signaling through the MAP kinase (MAPK) and PI-3 kinase (PI3K) pathways, which we measured by phosphorylation of Erk1/2 and Akt, respectively, was also normal (Fig. S2b). Conversely, antibody treatment dampened cytosolic calcium (Ca^2+^) influx (Fig. S2c) and inhibited the upregulation of CD69, an immediate early response gene (Fig. 1f and S1f). To decouple these LFA-1 dependent effects on T cell activation from a distinct and specific role in degranulation, we used a combination of phorbol myristate acetate (PMA) and the Ca^2+^ ionophore A23187 (Iono) to induce T cell activation in the absence of TCR engagement (Fig. 1c, S1c). CTLs treated with PMA/Iono alone exhibited robust CD69 expression (Fig. 1f, S1f), indicative of strong activation. Their degranulation responses were quite modest, however (Fig. 1d, S1e), pointing to the importance of target contact/proximity in stimulating cytotoxic secretion. Indeed, robust Lamp1 exposure was only observed in CTLs that were concomitantly exposed to antigenic pMHC and ICAM, either on target cells or on beads. Critically, this ligand-induced component of degranulation was completely inhibited by LFA-1 blockade (Fig. 1d, S1e). Taken together, these results indicate that LFA-1 engagement promotes cytotoxic secretion independently of T cell activation and at a level downstream of early signaling events. Notably, LFA-1 was also required for the formation of strong CTL-target cell conjugates, both in the presence and in the absence of PMA/Iono (Fig. 1g). Hence, the capacity of LFA-1 to stimulate degranulation was phenotypically linked to its ability to mediate strong adhesion.

That both the TCR and LFA-1 were required for robust degranulation (Fig. 1 and S1) raised the possibility that both receptor types must be engaged within the same cell-cell interface to elicit cytotoxic responses. This requirement would presumably enhance the specificity of killing by ensuring that toxic factors are released only against *bona fide* target cells (expressing cognate pMHC) that are tightly associated with the CTL (via integrin adhesion). To test this idea, we stimulated CTLs using beads coated with both pMHC and ICAM-1 (*cis*) or using a mixture of beads coated separately with only pMHC or ICAM-1 (*trans*) (Fig. 2a). Care was taken to make sure that the total amount of accessible pMHC and ICAM-1 was identical in each experimental group and that CTLs could engage multiple beads simultaneously (Fig. S3a). Strikingly, immobilized ICAM-1 boosted TCR-induced degranulation only when presented in the *cis* configuration (Fig. 2b). Indeed, CTLs stimulated with the *trans* mixture of pMHC- and ICAM-1-coated beads responded indistinguishably from CTLs stimulated with pMHC beads alone. We observed the same pattern of results using CD69 upregulation as the downstream readout (Fig. 2c). Hence, ligand-bound LFA-1 must share the same interface as the ligand-bound TCR in order to boost cytotoxicity and T cell activation.

**Figure 2.**
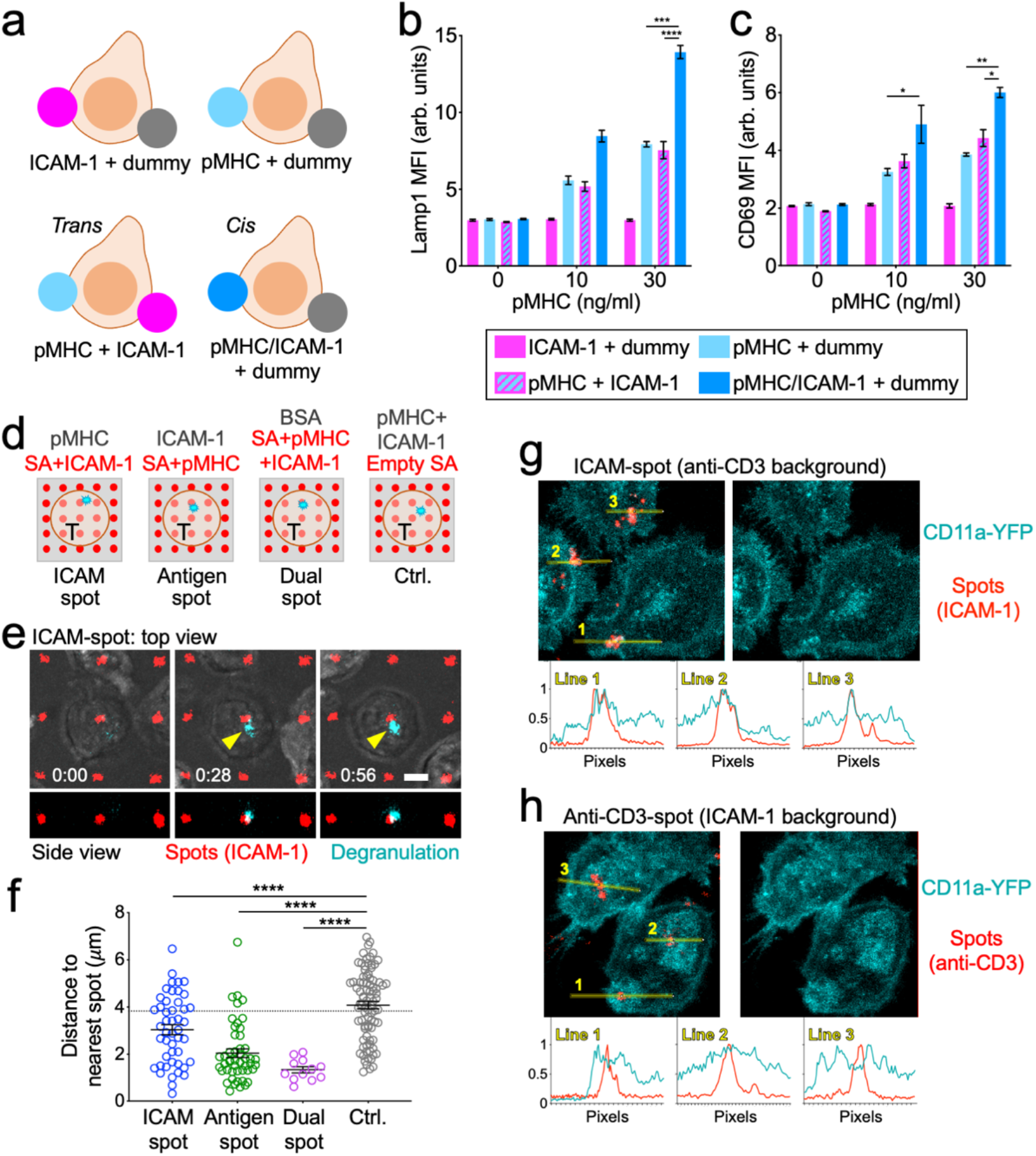
Degranulation occurs in IS domains containing both ligand-bound TCR and ligand-bound LFA-1. (a-c) OT-1 CTLs were activated by equal mixtures of stimulatory beads bearing the indicated proteins. (a) A schematic of the experiment. Dummy denotes beads coated with nonstimulatory pMHC (H-2D^b^-KAVY) alone. *Trans* and *cis* copresentation of H-2K^b^-OVA and ICAM-1 are indicated. (b-c) Graphs showing CTL degranulation (b) and CD69 expression (c), measured 90 min after CTL stimulation. Data were derived from technical triplicates. (d-f) OT-1 CTLs expressing pHluorin-Lamp1 were imaged on micropatterned surfaces coated with stimulatory pMHC (H-2K^b^-OVA) and ICAM-1 in 4 different configurations (d). SA = streptavidin. Time-lapse montage showing a representative degranulation event (indicated by the yellow arrowhead) on an ICAM-spot surface. In the top views, fluorescent signals have been superimposed on the brightfield image. Time in M:SS is shown at the bottom left corner of the top view images. Scale bars = 4 μm. (f) Distance between each degranulation event and the closest fluorescent SA spot in the IS. Dotted line indicates the mean distance expected from randomly placed degranulation. N = 12 for Dual-spot and ≥ 47 for the other conditions. (g-h) CD11a-YFP CTLs were imaged on ICAM-spot (g) and Anti-CD3-spot (h) surfaces. Representative images are shown in the top of each panel, with linescans through the fluorescent SA spots below. The lines used to generate the linescans are shown in the left image of each panel. All error bars denote SEM. *, **, ***, and **** denote P ≤ 0.05, P ≤ 0.01, P ≤ 0.001, and P ≤ 0.0001, calculated by 1way and 2way (b-c) ANOVA. All data are representative of at least two independent experiments.

Building upon this idea, we next examined whether local coengagement of LFA-1 and the TCR could control the position of degranulation events within the IS. Using protein microstamping^19^, we prepared glass coverslips containing 2 μm spots of fluorescent streptavidin spaced in a 10 μm × 10 μm square grid. These surfaces were then incubated with mixtures of unbiotinylated proteins to coat/block the empty glass between streptavidin spots, and then with biotinylated proteins to load the spots themselves. By varying the composition of coating and loading mixtures, we were able to generate a panel of distinct micropatterned substrates: 1) ICAM-1 spots within a uniform background of pMHC (ICAM-spot), 2) pMHC spots on an ICAM-1 background (Antigen-spot), and 3) spots containing both ICAM-1 and pMHC on a nonstimulatory (BSA coated) background (Dual-spot) (Fig. 2d). We also generated control surfaces containing empty streptavidin spots in a background of admixed pMHC and ICAM-1. CTLs plated on Dual-spot surfaces evinced Ca^2+^ flux only during periods of spot contact (Fig. S3b and Movies S1-2), confirming that biotinylated stimulatory ligands could be constrained in space by the patterned streptavidin. To monitor degranulation position in this system, we employed a reporter construct in which the pH-sensitive fluorescent protein pHluorin is linked to the lytic granule resident protein Lamp1^20^. Because pHluorin is quenched in the low pH granule environment, it becomes visible only upon fusion, generating a transient burst of fluorescence that reveals the position of the degranulation site within the IS. When CTLs expressing pHluorin-Lamp1 were imaged on ICAM-spot, Antigen-spot, or Dual-spot surfaces, degranulation events tended to cluster around the fluorescent spots containing ICAM-1 and/or pMHC (Fig. 2e-f and Movie S3). In all three cases, the mean distance between degranulation events and the spots closest to them was substantially lower than one would expect by chance (dotted line in Fig. 2f) and significantly less than the corresponding distances measured between empty SA spots and degranulations on control surfaces. The enrichment of degranulation in zones where pMHC and ICAM-1 were either copresented (Dual-spot) or closely apposed (ICAM-spot and Antigen-spot) further supports a critical role for the coengagement of the TCR and LFA-1 in guiding cytotoxic secretion and suggests that permissive secretory domains of receptor coengagement can be substantially smaller than the IS itself.

Although all three micropatterned substrates elicited targeted degranulation, responses to the Dual-spot and Antigen-spot configurations were significantly more focused than what we observed on ICAM-spot surfaces (Fig. 2f). To explore the basis for this difference, we imaged CTLs expressing a fluorescent form of the LFA-1 α-chain (CD11a-YFP) on surfaces containing focal LFA-1 or TCR ligands. Because the CD11a-YFP CTLs were derived from a polyclonal animal rather than an OT-1 transgenic, we used an anti-CD3 antibody instead of antigenic pMHC to engage the TCR. On ICAM-spot surfaces (anti-CD3 in the background), LFA-1 tended to localize to the micropatterned ICAM-1 (Fig. 2g), consistent with ligand recognition. The Anti-CD3-spot configuration, however, induced even stronger focal accumulation of LFA-1 (Fig. 2h), which was surprising considering that ICAM-1 was coated in the background of these surfaces. Interestingly, LFA-1 recruitment on Anti-CD3-spot substrates was most apparent not over the anti-CD3 spot itself but in the surrounding 1-2 μm neighborhood. This could potentially reflect inside-out integrin activation and clustering induced by local TCR signaling^15^. The unexpectedly robust accumulation of LFA-1 to areas of focal TCR stimulation potentially explains why Antigen-spot surfaces elicited more focused degranulation than their ICAM-spot counterparts.

Within the IS, both the TCR and LFA-1 are subjected to F-actin dependent pulling forces, which are thought to drive the formation of catch bonds between each receptor and its respective ligand, promote conformational changes, and induce signal transduction^10, 11, 14, 21^. Given the importance of these forces for the function of each receptor, we reasoned that they might also play a role in guiding cytotoxic secretion. To investigate this hypothesis, we employed Förster resonance energy transfer (FRET)-based molecular tension probes (MTPs)^22^ specific for the TCR and LFA-1. Each MTP comprised a stimulatory ligand (pMHC or ICAM1) attached to a DNA hairpin containing a fluorophore at one end (Atto647N or Cy3B, respectively) and a quencher (BHQ-2) at the other (Fig. 3a). When folded at resting state, MTPs do not fluoresce due to the close proximity between quencher and fluorophore. Applied forces capable of unwinding the hairpin (in this case, 4.7 pN) pull the quencher and fluorophore apart, dramatically increasing fluorescence. Consistent with prior reports^23, 24^, surfaces coated with pMHC-MTPs and ICAM-1-MTPs induced IS formation by OT1 CTLs and the exertion of dynamic forces through both LFA-1 and the TCR (Fig. 3b and Movie S4), which we visualized by time-lapse imaging. To measure the association between degranulation and receptor-specific forces, we used CTLs expressing pHluorin-Lamp1 to record exocytic events elicited by stimulatory MTPs (Fig. 3c). The mean MTP fluorescence in the immediate vicinity of each event (2 μm box) was then compared with the mean fluorescence of the entire IS. This approach revealed a marked enrichment of ICAM-1-MTP signal in the degranulation zone (Fig. 3d), indicative of a spatial correlation between cytotoxic secretion and force exertion through LFA-1. pMHC-MTP pulling was not associated with degranulation in this way (Fig. 3d), arguing against a role for the TCR as a critical force bearing receptor in this context. To further characterize the pattern of LFA-1 mechanics, we examined ICAM-1-MTP fluorescence along linescans bisecting the degranulation peak. Mean LFA-1 forces reached a local maximum in the 1 μm diameter region surrounding each event (Fig. 3e), consistent with the idea that mechanically active LFA-1 defines permissive zones for cytotoxic secretion. A degranulation zone of this size would accommodate the approach of a typical lytic granule (0.5 – 1μm in diameter)^25^.

**Figure 3.**
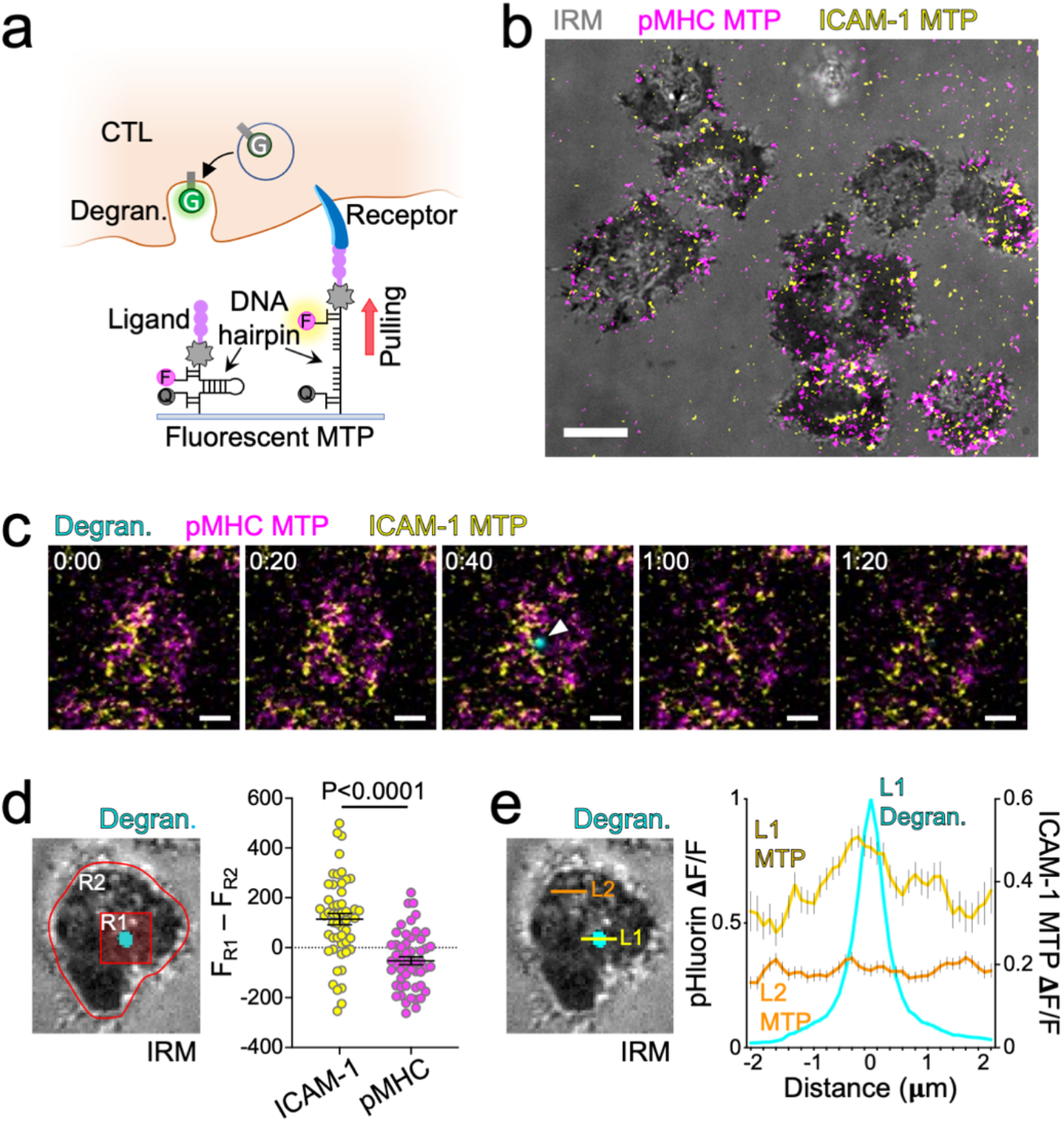
LFA-1 pulling forces define degranulation domains. (a) Measuring correlations between degranulation (Degran.) and receptor specific pulling forces with MTPs. F = fluorophore, Q = quencher, G = pHluorin. (b-e) OT-1 CTLs expressing pHluorin-Lamp1 were imaged by TIRF microscopy on glass surfaces coated with pMHC (H-2K^b^-OVA) and ICAM-1 MTPs. (b) Representative image of pMHC-MTP and ICAM-1-MTP signals, overlaid onto the corresponding IRM image. Scale bar = 5 μm. (c) Time-lapse montage showing a representative degranulation event (indicated by a white arrowhead) together with pMHC-MTP and ICAM-1-MTP signals. Time in M:SS is shown at the top left corner of each image. Scale bars = 2 μm. (d) Left, image of a representative degranulation event, overlaid onto the corresponding IRM image. Regions defining the degranulation subdomain (R1) and the entire IS (R2) are indicated. Right, differences in mean fluorescence intensity between R1 and R2 at the moment of degranulation are shown for the indicated MTPs. N = 52 for each sample. Error bars denote SEM. P value calculated by unpaired Student’s t test. (e) Left, image of a representative degranulation event, overlaid onto the corresponding IRM image. Linescans sampling the degranulating (L1) and inactive (L2) domains are indicated. Right, normalized ICAM-1-MTP fluorescence along L1 and L2 at the moment of degranulation (see Methods). The pHluorin-Lamp1 (Degran.) signal along L1 is shown for reference. Error bars denote SEM. N = 122 cells, pooled from two independent experiments. All other data are representative of at least two independent experiments.

Integrins are coupled to the F-actin cytoskeleton via talin, a mechanosensitive scaffolding protein that is critical for the formation and signaling of integrin adhesions^26^ (Fig. 4a). To evaluate the importance of integrin-cytoskeletal linkage for synaptic force exertion, we used CRISPR/Cas9 to deplete talin from OT1 CTLs (Fig. S4a) and then compared the physical output of these cells to that of controls expressing a nontargeting guide RNA. Talin depletion strongly suppressed ICAM-1 pulling on MTP surfaces (Fig. 4b-c and Movie S5), indicative of a profound defect in LFA-1 dependent force exertion. By contrast, pMHC-MTP forces were essentially unchanged (Fig. 4b-c and Movie S6), indicating that the mechanical effects of talin were restricted to LFA-1 in this system. The selectivity of the talin loss-of-function phenotype allowed us to interrogate the specific role of integrin mechanotransduction in cytotoxicity assays. In cocultures with OVA-loaded RMA-s cells, CTLs lacking talin exhibited sharply reduced degranulation and target cell lysis (Fig. 4d-e), implying a central role for integrin adhesions in both processes. These loss-of-function phenotypes were not rescued by the application of PMA/Iono (Fig. 4d-e), indicating that they were not caused by impaired T cell activation. Consistent with this interpretation, depletion of talin did not affect TCR-induced MAPK and PI3K signaling (Fig. S4b), and it only modestly suppressed CD69 responses (Fig. 4f). The disproportionately large effect of talin deficiency on cytotoxic secretion was particularly obvious in two-dimensional plots of Lamp1 and CD69 (Fig. S4c), which confirmed that, for a given level of activation, CTLs lacking talin consistently degranulated more weakly than nontargeting controls. Talin depletion also suppressed CTL-target cell conjugate formation (Fig. 4g), similar to the effects of LFA-1 blockade (Fig. 1g). Collectively, these results support a critical role for talin in LFA-1 dependent IS mechanics and strongly suggest that it is force exertion through LFA-1, rather than the TCR, that imposes spatiotemporal control over CTL degranulation.

**Figure 4.**
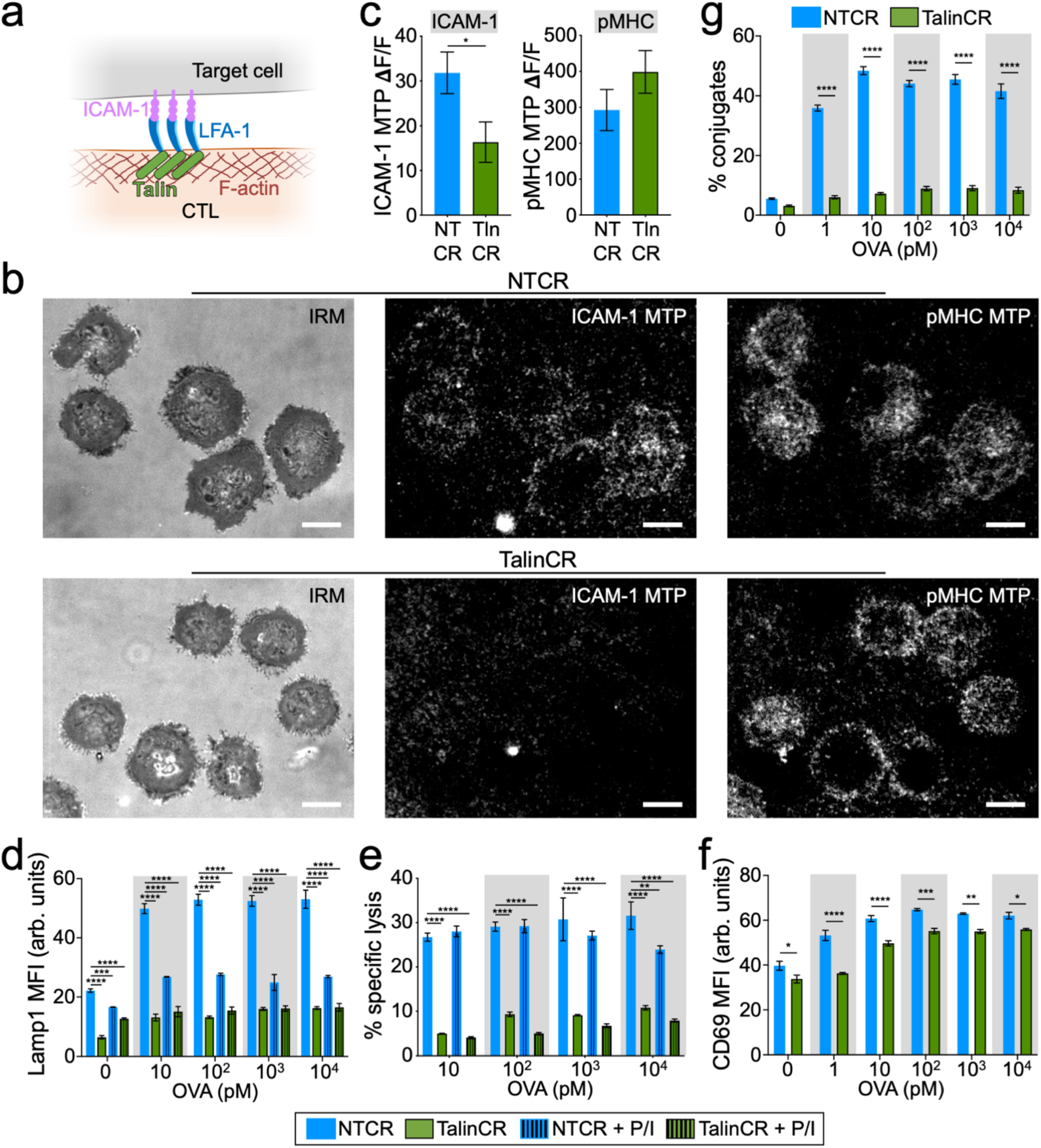
Talin is required for LFA-1 mediated force exertion, degranulation, and cytotoxicity. (a) Talin couples ligand-bound integrins to the F-actin cytoskeleton. (b-c) OT-1 Cas9 CTLs expressing talin gRNA (TalinCR) or control nontargeting gRNA (NTCR) were imaged on surfaces bearing the indicated stimulatory MTPs. (b) Representative images showing ligand specific pulling in synapses. Scale bars = 5 μm. (c) Mean ICAM-1-MTP (left) and pMHC (H-2K^b^-OVA)-MTP (right) signals, assessed 30 min after the addition of CTLs to the MTP surface. N ≥ 14 cells for each sample. (d-g) OVA-loaded RMA-s target cells were mixed with OT-1 Cas9 CTLs expressing the indicated gRNAs. PMA/Iono was applied to some samples in order to drive TCR independent CTL activation. (d) Lamp1 exposure (degranulation), measured 90 min after CTL-target cell mixing. (e) Target cell killing, measured 4 h after CTL-target cell mixing. (f) CD69 expression, measured 90 min after CTL-target cell mixing. (g) Conjugate formation, measured 90 min after CTL-target cell mixing. Data in d-g were derived from technical triplicates. All error bars denote SEM. *, **, ***, and **** denote P ≤ 0.05, P ≤ 0.01, P ≤ 0.001, and P ≤ 0.0001, calculated by unpaired Student’s t test (c) and 2way ANOVA (d-g). All data are representative of at least two independent experiments.

Not all cells express LFA-1 ligands, raising the question of whether integrin mechanotransduction controls CTL-mediated killing across a broad spectrum of targets. The capacity of talin deficiency to interrogate integrin function independently of LFA-1 allowed us to address this issue. B16F10 melanoma cells do not express ICAM-1, implying that they cannot engage LFA-1 across the IS (Fig. 5a-b). Nevertheless, they are reasonable targets for OT-1 CTLs, eliciting robust degranulation and cytotoxicity responses in the presence of OVA. LFA-1 blockade failed to inhibit either of these responses (Fig. 5c), consistent with the idea that LFA-1 is not involved in the recognition and killing of B16F10 cells. By contrast, talin depletion abrogated both target cell lysis and degranulation (Fig. 5d), strongly suggesting that integrins other than LFA-1 contribute to the killing of ICAM deficient targets. We conclude that integrin-mediated control of cytotoxic secretion is likely to be a general feature of the IS.

**Figure 5.**
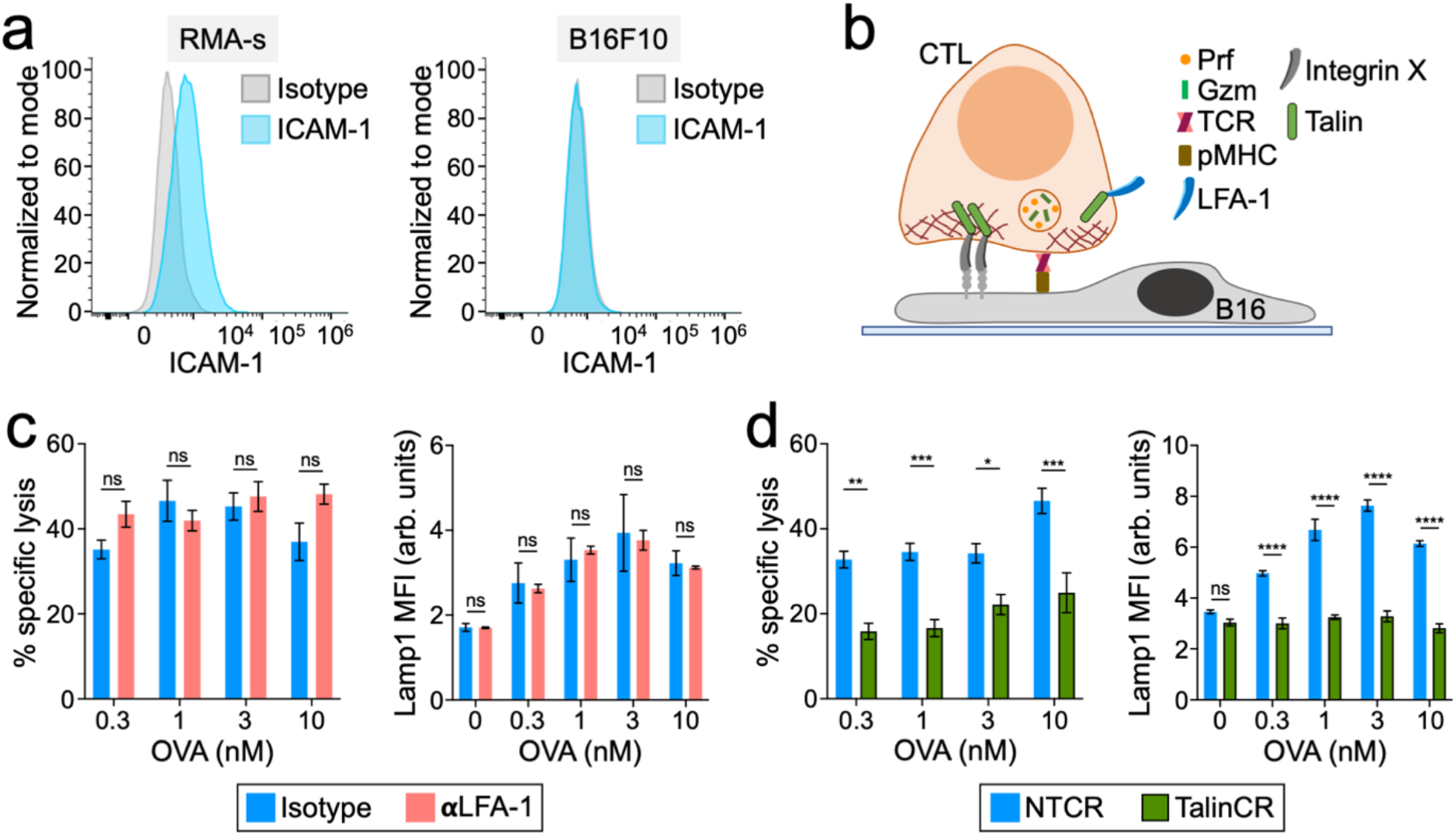
Talin, but not LFA-1, is required for CTL-mediated killing of B16F10 cells. (a) Representative histograms showing ICAM-1 expression in RMA-s (left) and B16F10 (right) cells. (b) Model of CTL-mediated killing of B16F10 cells that requires both the TCR and an unidentified integrin X. (c) OVA-loaded B16F10 target cells were mixed with OT-1 CTLs in the presence of LFA1 blocking antibody (αLFA-1) or isotype control. Left, specific lysis, measured after 4 h. Right, Lamp1 exposure, measured after 90 min. (d) OVA-loaded B16F10 target cells were mixed with OT-1 Cas9 CTLs transduced with the indicated gRNAs. Left, specific lysis, measured after 4 h. Right, Lamp1 exposure, measured after 90 min. Data in c-d were derived from technical triplicates. All error bars denote SEM. *, **, and *** denote P ≤ 0.05, P ≤ 0.01, and P ≤ 0.001, calculated by 2way ANOVA. All data are representative of at least two independent experiments.

Taken together, our data suggest a model in which degranulation occurs at permissive secretory subdomains within the IS that are defined by mechanically active integrins (Fig. S5). TCR signaling plays critical role in this process by inducing close contact formation and also by triggering Ca^2+^ influx, which is required for granule fusion^27-29^. In the absence of integrin-dependent force exertion, however, TCR signaling alone is insufficient for robust cytotoxic secretion. Indeed, our results imply that one of the major ways that the TCR promotes killing is by locally activating LFA-1 within the IS. This integrin-based model for degranulation both explains our data and is consistent with prior work documenting lytic granule accumulation and perforin release in synaptic subdomains defined by the engagement of adhesive and activating receptors^30, 31^. In the absence of LFA-1 ligands, we speculate that other integrins may assume its licensing role. Indeed, the fact that B16F10 cell killing requires talin, but not LFA-1, strongly implies the existence of alternative integrin activators. CTLs express both VLA-4 (α_4_β_1_) and CD103 (α_E_β_7_), which recognize protein ligands (VCAM-1/2 and E-Cadherin, respectively) found on subsets of potential target cells. CD103 is a particularly interesting candidate, as it has been shown to promote granule polarization and release toward E-Cadherin expressing tumor cells^32^.

Previous studies indicate that synaptic forces enhance the pore forming activity of perforin by straining the target membrane^8, 12^. The integrin dependent targeting model described above would be expected to facilitate this process by directing perforin to mechanically active subdomains within the IS. Integrin-mediated mechanotransduction also provides an elegant mechanism for identifying regions of synaptic membrane that are tightly engaged with the target cell, where extensive mechanical coupling between directly opposing membranes would enable the CTL to “feel” the presence of the target by pushing or pulling against it. Guiding degranulation to these regions of close apposition would ensure that only the target is exposed to perforin and granzyme, thereby limiting damage to innocent bystander cells. Hence, using mechanically active integrins to license cytotoxic secretion likely promotes both the potency and the specificity of killing responses.

The centrosome and its associated microtubules are key determinants of lytic granule localization^2, 4^. Target recognition induces the trafficking of lytic granules along microtubules toward the centrosome, which concomitantly reorients to a position just beneath the center of the IS, thereby positioning granules close to the synaptic membrane. Although this mechanism is thought to promote polarized secretion, studies from multiple labs indicate that it is dispensable for the process^5-7^. Indeed, we have found that CTLs lacking a functional centrosome or even the entire microtubule cytoskeleton retain the capacity to release perforin and granzyme directionally into the IS^5^. The integrin licensing model described here explains these prior observations by providing an alternative targeting mechanism. That being said, our results do not exclude an important role for the centrosome and microtubules in enhancing the speed and efficiency of cytotoxic secretion. Indeed, we have shown that microtubule depletion, while failing to disrupt the directionality of degranulation, nevertheless profoundly reduces the magnitude of the secretory response^5^. Accordingly, we favor a model in which centrosome polarization delivers granules into the IS neighborhood, at which point their site of fusion is dictated by integrin licensing.

The coupling of mechanical input to secretory output is unlikely to be unique to the cytotoxic IS. Indeed, one can imagine analogous mechanisms regulating other mechanically active processes, like phagocytosis and cell-cell fusion, that involve secretion and/or polarized membrane remodeling. We anticipate that biophysical analysis of systems like these will further illuminate the scope and functional relevance of mechano-secretory crosstalk in communicative cell-cell interactions.

## Methods

### Constructs

The retroviral expression construct for pHluorin-Lamp1 has been described^20^. A CRISPR gRNA construct targeting talin was prepared according to a published protocol^33^ using the following targeting sequence: 5’-GCTTGGCTTGTGAGGCCAGT-3’. A non-targeting control construct was also prepared using the sequence 5′-GCGAGGTATTCGGCTCCGCG-3′. After PCR amplification, DNA fragments encoding these guide sequences were subcloned into the pMRIG vector using the BamHI and MfeI restriction sites.

### Proteins

Class I MHC proteins (H-2K^b^ and H-2D^b^) were overexpressed in E. Coli, purified as inclusion bodies, and refolded by rapid dilution in the presence of β2-microglobulin and either OVA (for H-2K^b^) or KAVYDFATL (KAVY, for H-2D^b^). Monomeric MHC proteins were biotinylated using the BirA enzyme and purified by size exclusion chromatography. The extracellular domain of mouse ICAM-1 (a.a. 28-485, polyhistidine-tagged) was expressed by baculoviral infection of Hi-5 cells and purified by Ni^2+^ chromatography. After BirA-mediated biotinylation, the ICAM-1 was further purified by size exclusion.

### Cells

The animal protocols used for this study were approved by the Institutional Animal Care and Use Committee of Memorial Sloan Kettering Cancer Center. Primary CTL blasts were prepared by pulsing splenocytes from an OT1 αβTCR transgenic mice with 100 nM OVA in RPMI medium containing 10% (vol/vol) FCS. Cells were supplemented with 30 IU/mL IL-2 after 24 h and were split as needed in RPMI medium containing 10% (vol/vol) FCS and IL-2. RMA-s cells were maintained in RPMI containing 10% (vol/vol) FCS. B16F10 cells were maintained in DMEM medium containing 10% (vol/vol) FCS.

### Traction force microscopy

Arrays of PDMS (Sylgard 184; Dow Corning) micropillars (0.7 μm in diameter, 6 μm in height, spaced hexagonally with a 2-μm center-to-center distance) were cast onto glass coverslips using the inverse PDMS mold method^34^. After an ethanol wash and stepwise exchange into phosphate-buffered saline (PBS), pillars were stained with fluorescently labeled streptavidin (20 μg/mL Alexa Fluor 647, Thermo Fisher Scientific) for 2 hours at room temperature. Following additional PBS washes, the arrays were incubated with biotinylated H-2K^b^-OVA and ICAM-1 (10 μg/mL each) overnight at 4°C. The pillars were then washed into RPMI containing 5% (v/v) FCS and lacking phenol red for imaging. T cells stained with Alexa Fluor 488-labeled anti-CD45.2 Fab (clone 104-2) were then added to the arrays and imaged using an inverted fluorescence microscope (Olympus IX-81) fitted with a 100× objective lens and a mercury lamp for excitation. Images in the 488-nm (CTLs) and 647-nm (pillars) channels were collected every 15 s using MetaMorph software.

### Antibody blockade and pharmacological activation/inhibition

To assess the importance of LFA-1, CTLs were pre-incubated with LFA-1 blocking antibody (20 μg/mL, Clone M17/4, BioXCell) or an IgG2aκ isotype control antibody (20 μg/mL, Clone RTK2758, BioLegend) at 37ºC for 5 minutes before the addition of target cells/stimulatory beads. The final concentration of antibodies during the assay was 10 μg/mL. To induce T cell activation independently of the TCR, CTLs were pre-incubated with phorbol myristate acetate (PMA, 20 ng/mL, Sigma Aldrich) and the Ca^2+^ ionophore A23187 (2 μM, Tocris Bioscience) at 37 ºC for 5 minutes before the addition of target cells/stimulatory beads, yielding final concentrations of 10 ng/mL PMA and 1 μM A23187.

### Retroviral transduction

Phoenix E cells were transfected with expression vectors and packaging plasmids using the calcium phosphate method. Ecotropic viral supernatants were collected after 48 h at 37 °C and added to 1.5 × 10^6^ OT-1 blasts 24 h after primary peptide stimulation. Mixtures were centrifuged at 1400 × g in the presence of polybrene (4 μg/mL) at 35°C, after which the cells were split 1:3 in RPMI medium containing 10% (vol/vol) FCS and 30 IU/mL IL-2 and allowed to grow for an additional 4-6 days.

### Functional assays

To measure cytotoxicity, RMA-s target cells were labeled with CellTrace Violet (CTV), loaded with increasing concentrations of OVA, and mixed 3:1 with PKH26-stained OT-1 CTLs in a 96-well V-bottomed plate. Specific lysis of CTV^+^ target cells was determined by flow cytometry after 4 h at 37 °C^35^. To quantify degranulation, OT-1 CTLs were mixed with RMA-s target cells as described above and incubated at 37 °C for 90 min in the presence of eFluor 660–labeled anti-Lamp1 (clone eBio1D4B; eBioscience). Cells were then stained with FITC-labeled anti-CD69 (clone) and subjected to flow cytometric analysis to quantify Lamp1 and CD69 staining. To measure conjugate formation, labeled OT-1 CTLs and RMA-s targets were mixed 1:1, lightly centrifuged (100 × g) to encourage cell contact, and incubated 20 min at 37 °C. Cells were then resuspended in the presence of 2% paraformaldehyde, washed in fluorescence-activated cell sorting buffer (PBS + 4% FCS), and analyzed by flow cytometry. Conjugate formation was quantified as (PKH26^+^CTV^+^)/(PKH26^+^). For B16F10 killing assays, B16F10 targets were cultured overnight on fibronectin and then pulsed with varying concentrations of OVA for 2 hours. OT1 CTLs were added at an 8:1 E:T ratio and incubated for 3-4 hours at 37 °C in RPMI medium supplemented with IL-2 (30 IU/mL). Target cell death was quantified with an LDH (lactate dehydrogenase) cytotoxicity assay kit (Clontech) using the manufacturer’s recommended protocol. To measure intracellular granzyme B depletion, OT1 CTLs were mixed 1:3 with OVA-loaded RMA-s cells and incubated for 4-6 hours at 37 °C. Intracellular granzyme B levels were then measured by flow cytometry after fixation, permeabilization, and staining with Alexa 647 labeled anti-granzyme B (clone GRB11, Biolegend. All functional assays were performed in triplicate. To quantify ICAM-1 expression, RMA-s or B16F10 cells were stained with a fluorescently labeled anti-ICAM-1 antibody CD54 (Clone YN1/1.7.4, BioLegend) or isotype control antibody (Clone RTK4530, Biolegend).

### CTL activation with stimulatory beads

Streptavidin-conjugated polystyrene beads (Spherotech) were coated with 1 μg/mL biotinylated ICAM-1 and/or various concentrations of biotinylated H-2K^b^-OVA. Nonstimulatory pMHC (H-2D^b^) was used if necessary to adjust the total biotinylated protein concentration of each mixture 2 μg/mL. After overnight incubation at 4°C, excess unbound protein was washed out and the beads were transferred into RPMI medium containing 10% (vol/vol) FCS (+/-phenol red) for use in experiments. For immunoblot analysis of signaling, beads were mixed with OT-1 CTLs at a 1:1 ratio. For functional studies (e.g. degranulation), the CTL to bead ratio was 1:3. For 2 bead stimulation (Fig. 2a-c), Nile Red and Purple streptavidin beads were coated as described above with H-2K^b^-OVA and ICAM-1 in either *cis* or *trans* configurations. H-2D^b^ was used to fill empty spaces in the *trans* configuration and also to generate “dummy” coated beads. In each experimental condition, CTLs were mixed with two kinds of beads at a 1:3:3 ratio. Degranulation and CD69 upregulation were quantified by flow cytometry as described above.

### Micropatterning experiments

PDMS stamps for imprinting 2 μm in diameter spots in 10 μm center-to-center square arrays were prepared using microfabricated silicon masters as previously described^19^. Stamps were washed in ethanol and water, then coated with fluorescently-labeled streptavidin (10 μg/mL Alexa Fluor 647, Thermo Fisher Scientific) for 1 hour at room temperature. After PBS washing to remove excess proteins, the stamps were pressed onto 35 mm glass coverslips (#1.5) to transfer the streptavidin. Coverslips were then incubated with the following unbiotinylated proteins to coat the spaces between streptavidin dots: (1) ICAM-spot – 10 μg/mL H-2K^b^-OVA, (2) Antigen-spot – 10 μg/mL ICAM-1, (3) Dual-spot – 5% BSA, (4) ICAM-spot with anti-CD3 background – 10 ug/mL anti-CD3 antibody (Clone 145-2C11, eBioScience), (5) Anti-CD3-spot – 10 μg/mL ICAM-1, and (6) control – 10 μg/mL unlabeled streptavidin. After 1 h at room temperature, coverslips were rinsed with PBS 3 times to wash away uncoated protein and blocked with 5% BSA at room temperature for 1 hour before another round of PBS washes. Coverslips were then incubated with the following biotinylated proteins to load the streptavidin: (1) ICAM-spot – 2 μg/mL ICAM-1, (2) Antigen-spot – 2 μg/mL H-2K^b^-OVA, (3) Dual-spot – 2 μg/mL H-2K^b^-OVA and 2 μg/mL ICAM-1, (4) ICAM-spot with anti-CD3 background – 2 μg/mL ICAM-1, (5) Anti-CD3-spot – 2 ug/mL anti-CD3 antibody (Clone 145-2C11, eBioScience), and (6) control – 2 μg/mL H-2K^b^-OVA and 2 μg/mL ICAM-1. After 1 h at room temperature, the surfaces were washed into RPMI containing 5% (v/v) fetal calf serum (FCS) and lacking phenol red for imaging. Cells were then added and imaging performed using either a Leica SP5-inverted confocal laser scanning microscope fitted with 488 nm, 563 nm, and 647 nm lasers, or a Leica SP8-inverted confocal laser scanning microscope fitted with a white light laser. In general, samples were imaged every 15 s for 30 min.

### Calcium imaging

CTLs were loaded with 5 μg/mL Fura2-AM (ThermoFisher Scientific), washed, and then imaged on stimulatory glass surfaces coated with H-2K^b^-OVA and ICAM-1 as previously described^36^. 340 nm and 380 nm excitation images were acquired every 30 seconds for 30 min using a 20× objective lens (Olympus).

### DNA hairpins

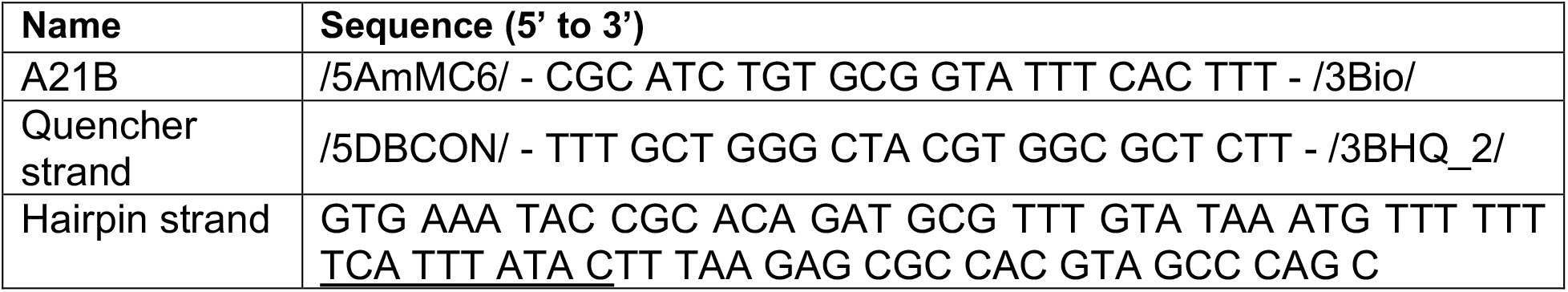

A mixture of oligo A21B (10 nmol) and excess Cy3B-NHS ester or Atto647N-NHS ester (50 μg) in 0.1 M sodium bicarbonate solution was allowed to react at room temperature overnight. The derivatized oligo was then purified by gel filtration and reversed phase HPLC.

### MTP surface preparation and imaging

Glass coverslips (No. 1.5H, Ibidi) were sonicated in MilliQ H_2_O and ethanol, rinsed in H_2_O, and then immersed in piranha solution (3:1 sulfuric acid:H_2_O_2_) for 30 min to remove organic residues and activate hydroxyl groups on the glass. Subsequently, the cleaned substrates were rinsed with more H_2_O and ethanol and then transferred to a 200 mL beaker containing 3% APTES in ethanol for 1 h, washed with ethanol and baked at ∼100 ºC for 30 min. After cooling, slides were mounted to 6-channel microfluidic cells (Sticky-Slide VI 0.4, ibidi). To each channel, ∼50 μL of 10 mg/mL of NHS-PEG4-azide in 0.1 M NaHCO_3_ (pH = 9) was added and incubated for 1 h. The channels were then washed with H_2_O, blocked with 0.1% BSA in PBS for 30 min, and washed with PBS. ∼50 μL of PBS solution was retained inside the channel after washing to prevent drying. Subsequently, the hairpin tension probes were assembled in 1M NaCl by mixing the Atto647N labeled A21B strand (220 nM), quencher strand (220 nM) and hairpin strand (200 nM) in the ratio of 1.1: 1.1:1. The mixture was heat annealed by incubating at 95 °C for 5 min, followed by cool down to 25 ºC over 30 min. ∼50 μL of the assembled probe was added to the channels (total volume = ∼100 μL) and incubated overnight at room temperature. The following day, unbound DNA probes were removed by PBS wash. Then, 10 μg/mL of streptavidin was incubated in the channels for 45 min at room temperature. The surfaces were cleaned with PBS and incubated with 5 μg/mL of biotinylated pMHC ligand for 45 min at room temperature. After PBS washing, a second DNA tension probe (Cy3B labeled) was assembled and attached as described above, followed by loading with streptavidin and 5 μg/mL of biotinylated ICAM-1. Monomeric ICAM-1 was used for pHluorin-Lamp1 experiments (Fig. 3) and dimeric Fc-ICAM1 for talin-KO experiments (Fig. 4). After washing off the unbound ICAM-1 protein, surfaces were rinsed in complete RPMI (no phenol red, supplemented with IL-2) in preparation for imaging with CTLs. MTP imaging was performed on a Nikon Eclipse Ti microscope attached to an electron multiplying charge coupled device (EMCCD; Photometrics), an Intensilight epifluorescence source (Nikon), a CFI Apo 100× (NA 1.49) objective lens (Nikon), and a TIRF launcher with 488 nm, 561 nm, and 638 nm laser lines. In general, IRM, 488 nm, 561 nm, and 638 nm images were collected every 20 s for 30 min. TIRF illumination was used to image pHluorin-Lamp1 and epifluorescence to image the MTPs.

### Imaging analysis

Imaging data were analyzed using SlideBook (3i), Imaris (Bitplane), Excel (Microsoft), Prism (GraphPad), and Python in Jupyter Notebook^37^. Ca^2+^ signaling was quantified by determining the mean Fura2 ratio for all cells in the imaging field using a mask thresholded on the 340 nm excitation signal. To quantify force exertion in traction force microscopy experiments, custom MATLAB scripts were used to extract pillar displacements from the imaging data, which were then converted into force vectors^16^. To measure the distance between degranulation events and the closest streptavidin spot on micropatterned surfaces, pHluorin-Lamp1 and streptavidin Alexa Fluor 647 signals were converted into Imaris spot constructs using Imaris scripts. The distances (μm) between each pHluorin-Lamp1 signal of interest and the closest streptavidin spot within the synaptic boundary of the CTL were then determined using the Imaris ‘Shortest Distance’ function. The expected distance between randomly placed degranulation events and ligand spots was determined in silico. First, the unit cell of the micropattern was modeled as a 5 μm × 5 μm square with a quarter-circle representing the stamped protein at one corner. Then, the unit square was divided into 1 × 10^6^ points (evenly sampling 10-nm spaces in both x and y), and the Euclidean distance of each point to the stamped protein corner was calculated. The mean distance of this distribution is 3.8266 μm. MTP data were analyzed by comparing the mean fluorescence intensity of each MTP within the 2 μm × 2 μm box centered on a pHluorin-Lamp1 signal of interest with the mean fluorescence intensity of the MTP within the entire IS, defined by threshold masking of IRM images (Fig. 3d). Linescan analysis of ICAM-1-MTP fluorescence at degranulation sites (Fig. 3e) was performed by generating a series of 2 μm linescans bisecting degranulation events of interest. The linescan intensities were aligned around the degranulation, averaged over each pixel, and normalized per linescan. An analogous set of control linescans, collected from parts of the IS lacking degranulation events, were processed in parallel using the same scripts.

### Proliferation Assay

Day 7 OT-1 CTLs were stained with CellTrace Violet at room temperature for 20 min, washed in serum containing medium, and then incubated with irradiated OVA-loaded C57BL/6 splenocytes (0.5 × 10^6^ CTL with 4.0 × 10^6^ splenocytes) the presence of 10 μg/mL anti-LFA-1 or an isotype control antibody. Subsequent dilution of CellTrace Violet was assessed by flow cytometry.

### Immunoblot

0.2-1 × 10^6^ CTLs were lysed using cold cell lysis buffer containing 50 mM TrisHCl, 0.15 M NaCl, 1 mM EDTA, 1% NP-40 and 0.25% sodium deoxycholate. Suppression of talin 1 was confirmed using an anti-talin 1 antibody (clone 8D4, Abcam). Actin served as a loading control (clone AC-15, Sigma). For signaling assays, serum and IL-2 starved OT1 CTLs were incubated with streptavidin polystyrene beads (Spherotech) coated with H-2K^b^-OVA and ICAM-1 at a 1:1 ratio for various times at 37 ºC and immediately lysed in 2× cold lysis buffer containing phosphatase inhibitors (1 mM NaF and 0.1 mM Na_3_VO_4_) and protease inhibitors (cOmplete mini cocktail, EDTA-free, Roche). Activation of PI3K and MAP kinase signaling was assessed by immunoblot for pAkt (Phospho-Akt (Ser473) Ab; Cell Signaling Technology) and pErk1/2 (Phospho-Thr202/ Tyr204; clone D13.14.4E; Cell Signaling Technology).

## Acknowledgements

We thank L. Stafford and C. Jeronimo for technical support; C. Sachar for assistance with traction force microscopy; Y. Romin, E. Chan, M. Tipping, V. Boyko, and the MSKCC Molecular Cytology Core Facility for assistance with confocal imaging; and members of the M.H. and L.C.K. laboratories for advice. Supported in part by the NIH (R01-AI087644 to M.H. and P30-CA008748 to MSKCC) and the Ludwig Foundation for Cancer Immunotherapy (M. D. J.).

## Author contributions

M.S.W., Y. Hu., E.E.S., K.S., and M.H. designed the experiments.

M.S.W., Y. Hu., E.E.S., W.J., and M.H. collected the data.

M.S.W., Y. Hu., E.E.S., X.X., M.D.J., and M.H. analyzed the data.

N.H.R., J.H.L., Y. Hong, M.K., and L.C.K. contributed key reagents.

M.S.W. and M.H. wrote the paper.

---

**Figure S1.**
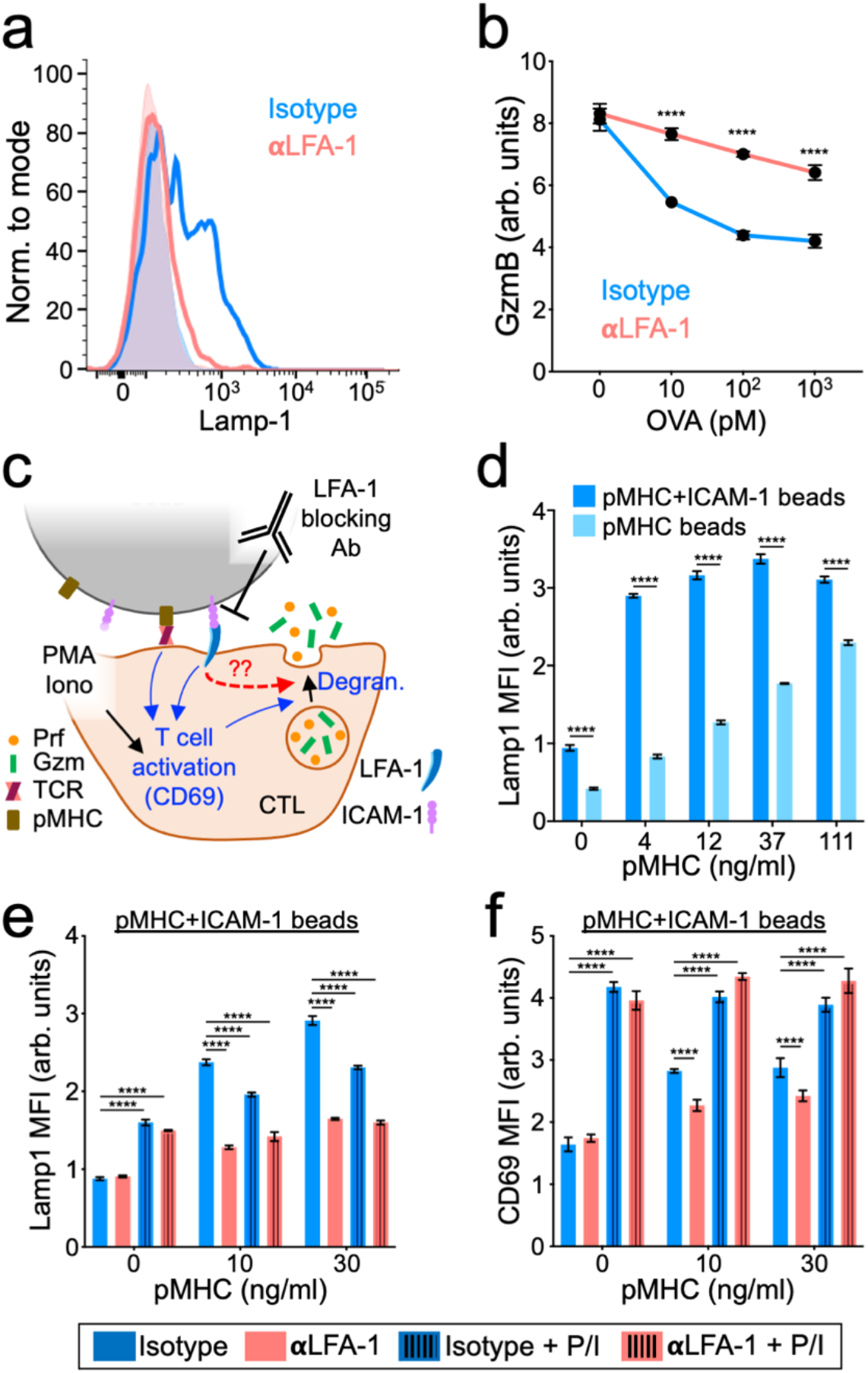
LFA-1 blockade disrupts CTL degranulation. (a-b) OT-1 CTLs were incubated with OVA-loaded RMA-s cells in the presence of LFA1 blocking antibody (αLFA-1) or isotype control. (a) A representative FACS histogram of CTL Lamp1 staining, measured 90 min after cell mixing. (b) Staining for intracellular granzyme B (GzmB) in CTLs, performed 90 min after cell mixing. (c) Diagram schematizing CTL activation by stimulatory beads coated with pMHC (H-2K^b^-OVA) ± ICAM-1. (d) OT-1 CTLs were mixed with beads coated with increasing amounts of pMHC in the presence or absence of ICAM-1. Degranulation was measured 90 min after CTL stimulation. (e-f) Beads coated with pMHC and ICAM-1 were incubated with OT-1 CTLs in the presence or absence of PMA/Iono and treated with either αLFA-1 or isotype control. Graphs show degranulation (e) and CD69 expression (f), measured 90 min after CTL stimulation. All error bars denote SEM. *, **, ***, and **** denote P ≤ 0.05, P ≤ 0.01, P ≤ 0.001, and P ≤ 0.0001, calculated by 2way ANOVA. All data are representative of at least two independent experiments.

**Figure S2.**
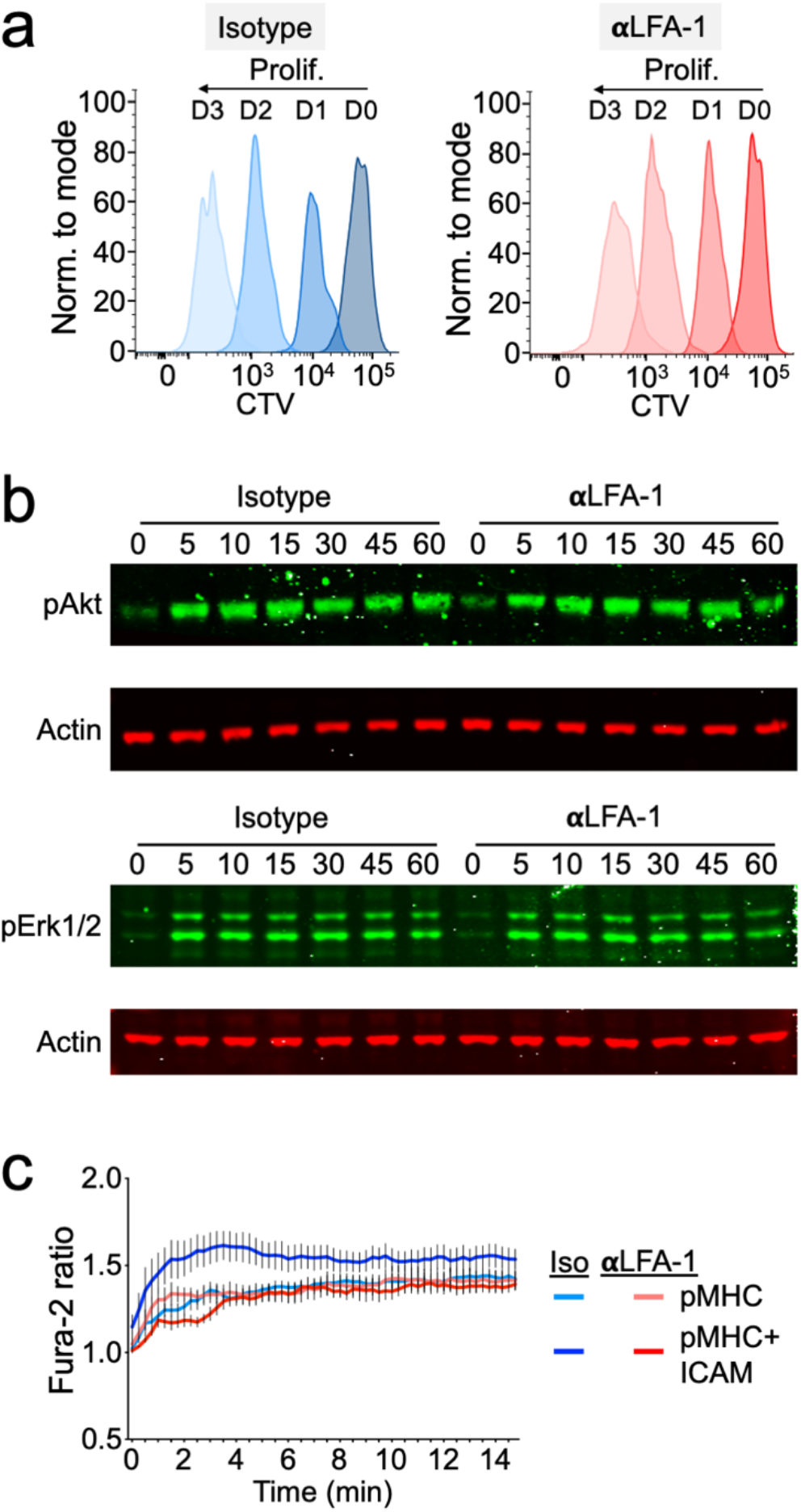
LFA-1 blockade alters only some indices of T cell activation. (a) OT-1 CTLs were labeled with CellTrace Violet (CTV) and incubated with OVA-loaded splenocytes in the presence of αLFA-1 or isotype control. CTV dilution was monitored over 3 days (D) by flow cytometry. (b) OT-1 CTLs were mixed with pMHC (H-2K^b^-OVA) and ICAM-1 coated beads and, at the indicated timepoints, pAKT (top) and pErk1/2 (bottom) were assessed by immunoblot, with actin serving as a loading control. (c) OT-1 CTLs were loaded with Fura-2-AM and imaged on glass surfaces coated with the indicated proteins in the presence of αLFA-1 or isotype control. Graph shows the mean Fura-2 ratio of all CTLs in the imaging field, averaged over 6 positions. Error bars denote SEM. All data are representative of at least two independent experiments.

**Figure S3.**
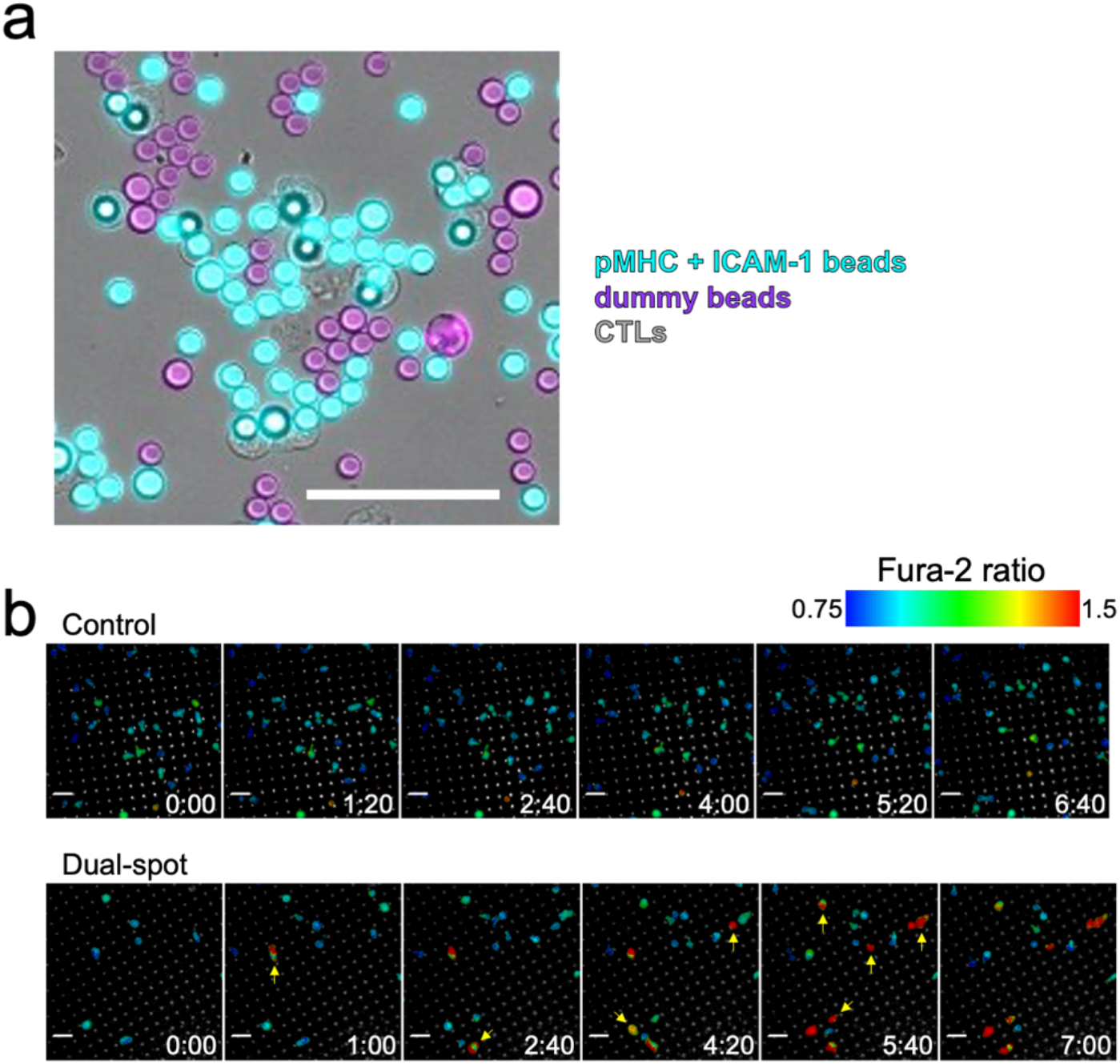
CTL stimulation with spatially segregated ligands. (a) Representative image of OT-1 CTLs, pMHC-coated beads, and ICAM-1 coated beads mixed at the proportions used for Figure 2a-c. Scale bar = 50 μm. (b) Representative time-lapse montages of Fura-2-AM-loaded OT-1 CTLs imaged on Dual-spot and control surfaces. Yellow arrows denote the onset of Ca^2+^ flux in individual cells. Time in M:SS is shown in the bottom right corner of each image. Scale bars = 10 μm. All data are representative of at least two independent experiments.

**Figure S4.**
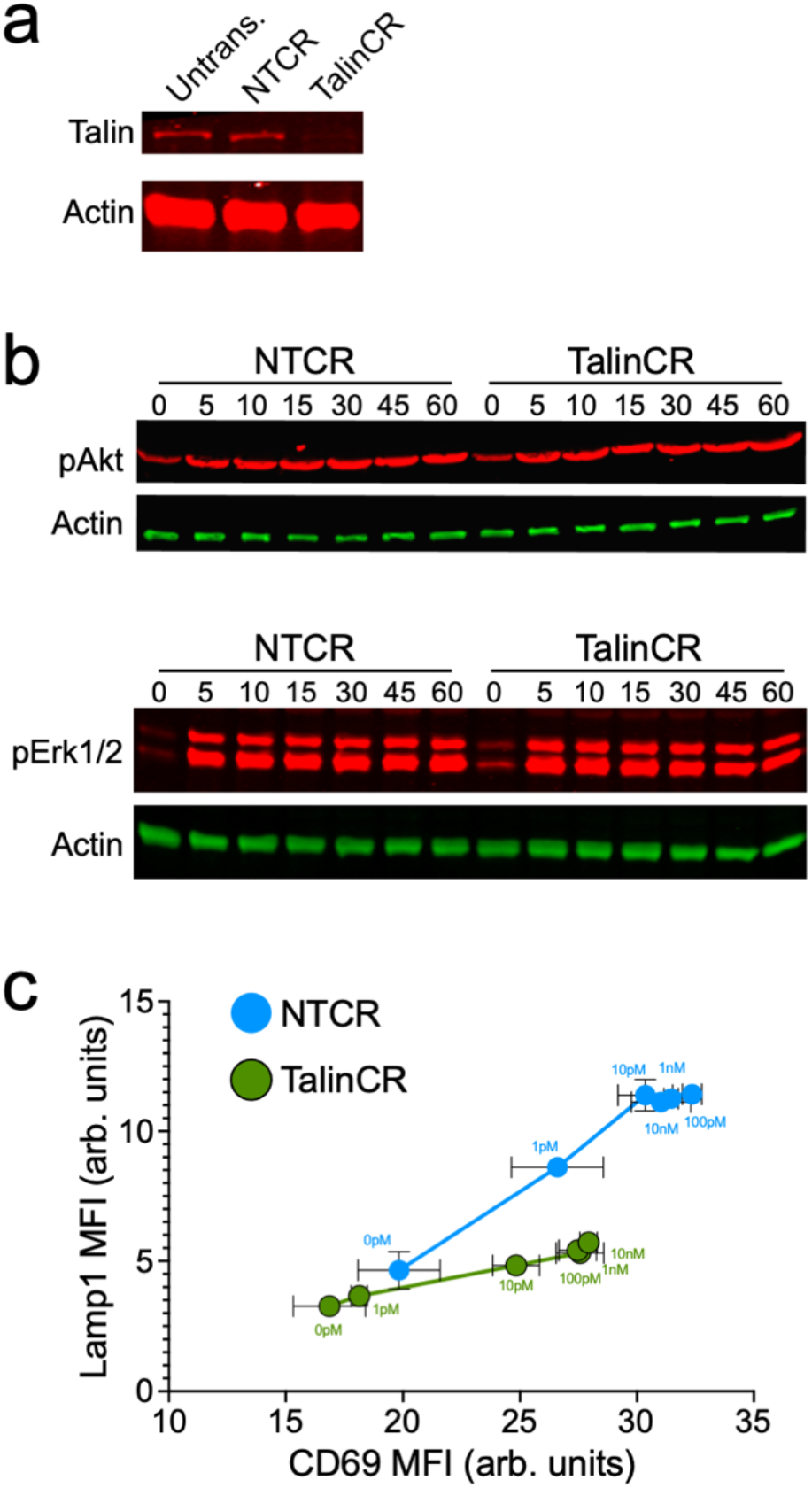
Talin depletion inhibits degranulation but not early TCR signaling. (a) Immunoblot analysis of talin expression in OT-1 Cas9 CTLs transduced with the indicated gRNAs. Untrans. = untransduced. (b) OT-1 Cas9 CTLs transduced with the indicated gRNAs were mixed with pMHC (H-2K^b^-OVA) and ICAM-1 coated beads and, at the indicated timepoints, pAKT (top) and pErk1/2 (bottom) were assessed by immunoblot. In a and b, actin served as a loading control. (c) 2-dimensional plot correlating degranulation (Lamp1) and T cell activation (CD69), taken from the same data set used to generate Fig. 4d and 4f. Error bars denote SEM. All data are representative of at least two independent experiments.

**Figure S5.**
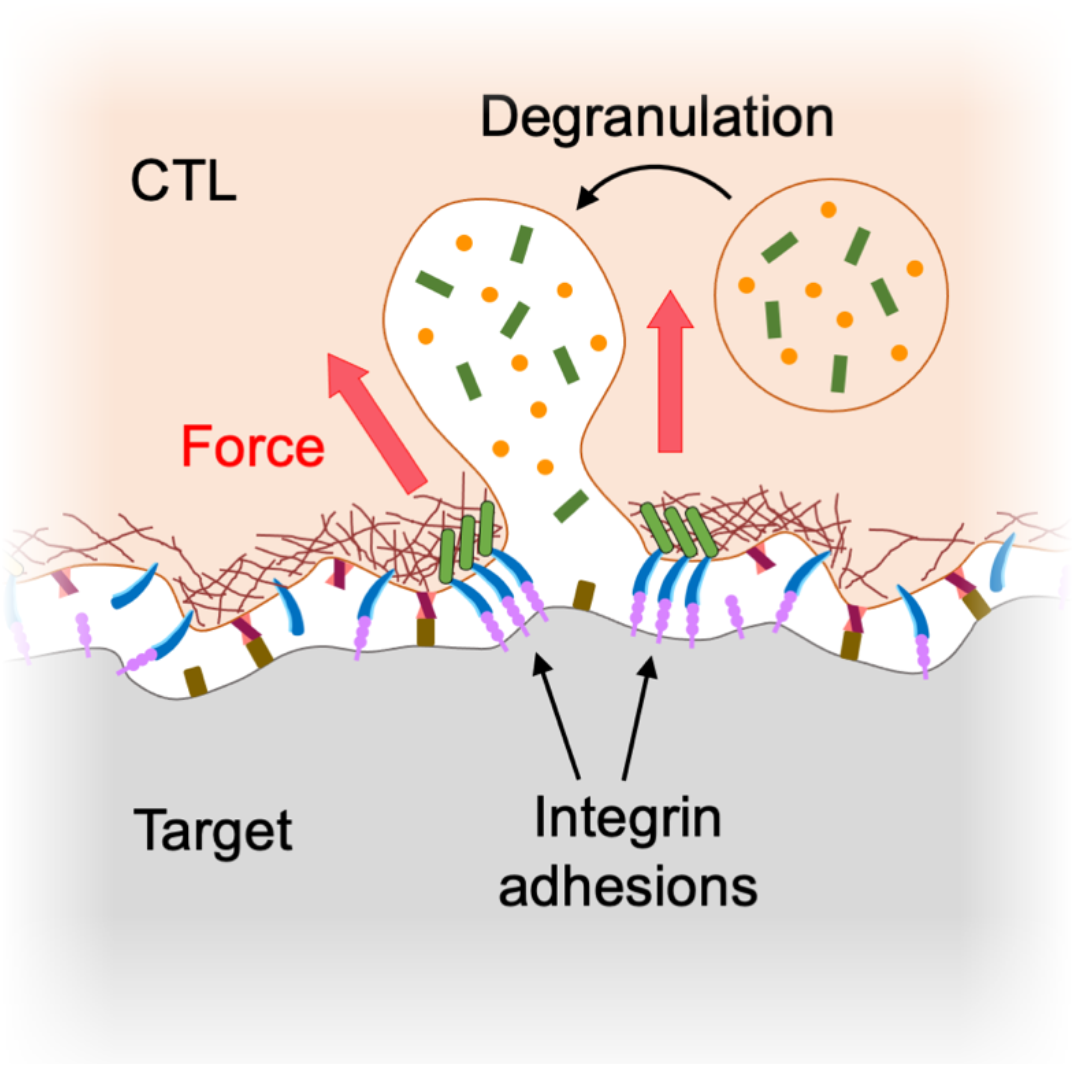
Mechanical licensing of degranulation by integrin adhesions. Lytic granules fuse within synaptic subdomains defined by ligand-bound integrins under tension.

## Supplementary Movie Legends

**Movie S1. Control micropatterned surfaces do not elicit T cell activation**. OT-1 CTLs loaded with Fura2-AM were imaged on control micropatterned surfaces containing empty fluorescent streptavidin dots. A representative 200× time-lapse movie is shown. Fura2 ratio is depicted in pseudocolor with cold and warm colors indicating low and high intracellular Ca^2+^, respectively. Time in HH:MM:SS is shown in the upper left corner. Scale bar = 10 μm.

**Movie S2. T cell activation on a stimulatory micropatterned surface**. OT-1 CTLs loaded with Fura2-AM were imaged on Dual-spot micropatterned surfaces with both pMHC and ICAM-1 loaded into fluorescent streptavidin dots. A representative 200× time-lapse movie is shown. Fura2 ratio is depicted in pseudocolor with cold and warm colors indicating low and high intracellular Ca^2+^, respectively. Time in HH:MM:SS is shown in the upper left corner. Scale bar = 10 μm.

**Movie S3. CTL degranulation on a micropatterned surface**. OT-1 CTLs expressing pHluorin-Lamp1 were imaged by confocal microscopy on ICAM-spot micropatterned surfaces. A representative 100× time-lapse movie is shown, with pHluorin-Lamp1 and Alexa Fluor 647 streptavidin depicted in cyan and red, respectively. Time in M:SS is shown in the bottom right. Scale bar = 2 μm. The CTL near the center of the field degranulates at 9:46.

**Movie S4. CTLs exert pulling forces through the TCR and LFA-1**. OT-1 CTLs were imaged on glass substrates coated with pMHC-Atto647N (cyan) and ICAM-1-Cy3B (yellow) MTPs. A representative 143× time-lapse movie is shown, with MTP fluorescence overlaid onto the corresponding IRM image. Time in MM:SS is indicated in the top left corner. Scale bar = 8 μm.

**Movie S5. Talin depletion inhibits force exertion through LFA-1, but not the TCR**. OT-1 Cas9 CTLs expressing talin specific gRNA (Talin CR) were imaged on glass substrates coated with pMHC-Atto647N and ICAM-1-Cy3B MTPs. Representative 143× time-lapse movies of MTP fluorescence are shown in montage together with the corresponding IRM signal. Time in MM:SS is indicated in the top left corner of the IRM time-lapse. Scale bar = 8 μm.

**Movie S6. Force exertion by talin sufficient CTLs**. OT-1 Cas9 CTLs expressing a nontargeting control gRNA (NT CR) were imaged on glass substrates coated with pMHC-Atto647N and ICAM-1-Cy3B MTPs. Representative 143× time-lapse movies of MTP fluorescence are shown in montage together with the corresponding IRM signal. Time in MM:SS is indicated in the top left corner of the IRM time-lapse. Scale bar = 8 μm.

